# Voluntary control of illusory contour formation

**DOI:** 10.1101/219279

**Authors:** William J Harrison, Reuben Rideaux

## Abstract

The extent to which visual inference is shaped by attentional goals is unclear. Voluntary attention may simply modulate the priority with which information is accessed by higher cognitive functions involved in perceptual decision making. Alternatively, voluntary attention may influence fundamental visual processes, such as those involved in segmenting an incoming retinal signal into a structured scene of coherent objects, thereby determining perceptual organisation. Here we tested whether the segmentation and integration of visual form can be determined by an observer’s goals by exploiting a novel variant of the classical Kanizsa figure. We generated predictions about the influence of attention with a machine classifier, and tested these predictions with a psychophysical response classification technique. Despite seeing the same image on each trial, observers’ perception of illusory spatial structure depended on their attentional goals. These attention-contingent illusory contours directly conflicted with equally plausible visual form implied by the geometry of the stimulus, revealing that attentional selection can determine the perceived layout of a fragmented scene. Attentional goals, therefore, not only select pre-computed features or regions of space for prioritised processing, but, under certain conditions, also greatly influence perceptual organisation and thus visual appearance.

**SIGNIFICANCE STATEMENT:** The extent to which higher cognitive functions can influence perceptual organisation is debated. The role of voluntary spatial attention, the ability to focus on only some parts of a scene, has been particularly controversial among neuroscientists and psychologists who aim to uncover the basic neural computations involved in grouping image features into coherent objects. To address this issue, we repeatedly presented the same novel ambiguous image to observers and changed their attentional goals by having them make fine spatial judgements about only some elements of the image. We found that observers’ attentional goals determine the perceived organisation of multiple illusory shapes. We thus reveal that voluntary spatial attention can control the fundamental processes that determine perceptual organisation.

## INTRODUCTION

The clutter inherent to natural visual environments means that goal-relevant objects often partially occlude one another. A critical function of the human visual system is to group common parts of objects while segmenting them from distracting objects and background, a process which requires interpreting an object’s borders. Figures which produce illusory contours, such as the classic Kanizsa triangle (Kanizsa, 1976), have provided many insights into this problem by revealing the inferential processes made in determining figure-ground relationships. These figures give rise to a vivid percept of a shape emerging from sparse information, and thus demonstrate the visual system’s ability to interpolate structure from fragmented information, to perceive edges in the absence of luminance discontinuities, and to fill-in a shape’s surface properties (for a review, see Shapley, Rubin, & Ringach, 2004). In the present study, we exploit these figures to investigate whether voluntary attention influences perceptual organisation.

Most objects can be differentiated from their backgrounds via a luminance-defined border. The visual system is tasked with allocating one side of the border to an occluding object, and the other side to the background. This computation can be performed by neurons in macaque visual area V2 whose receptive fields fall on the edge of an object (Zhou, Friedman, & von der Heydt, 2000). These “border-ownership” cells can distinguish figure from ground even when the monkey attends elsewhere in the display (Qiu, Sugihara, & von der Heydt, 2007), and psychophysical adaptation aftereffects suggest such cells also exist in humans (von der Heydt, Macuda, & Qiu, 2005). Further, neurophysiological work has revealed that V2 cells also process illusory edges (von der Heydt, Peterhans, & Baumgartner, 1984), though it is unclear whether those cells possess the same properties as border-ownership cells. These findings have contributed to the claim that visual structure is computed automatically and relatively early in the visual system, and that visual attention is guided by this pre-computed structure (Mihalas, Dong, von der Heydt, & Niebur, 2011).

It is also known, however, that visual attention can modulate the perception of figure-ground relationships of luminance-defined stimuli. Both voluntary and involuntary forms of attentional allocation can impact higher level cognition (Posner, 2014) and basic perception (Carrasco, 2011). Allocating visual attention to an area of space, for example, prioritises processing of stimuli presented at that location relative to other locations (Posner, 1980), and can alter apparent contrast (Carrasco, Ling, & Read, 2004). As early as 1832, Necker described his ability to alter the apparent depth of an engraved crystalline form, now referred to as a Necker cube, via an overt shift of attention (Necker, 1832). More recent psychophysical work has shown that voluntary attention can alter perceived depth order (Driver & Baylis, 1996) as in the case of Rubin’s face-vase illusion (Rubin, 1915)(Wagemans et al., 2012), surface transparency (Tse, 2005), speed (Anton-Erxleben, Henrich, & Treue, 2007; Turatto, Vescovi, & Valsecchi, 2007), contrast (Carrasco et al., 2004; Liu, Abrams, & Carrasco, 2009; Stormer, McDonald, & Hillyard, 2009) and spatial frequency (Abrams, Barbot, & Carrasco, 2010; Gobell & Carrasco, 2005). Furthermore, visual attention has been shown to facilitate visual grouping according to Gestalt rules at both the neurophysiological (Wannig, Stanisor, & Roelfsema, 2011) and behavioural (Barbot, Liu, Kimchi, & Carrasco, 2018; Houtkamp, Spekreijse, & Roelfsema, 2003) level. For instance, (Barbot et al., 2018) found that the apparent perceptual organization of luminance defined multielement arrays is either intensified or attenuated by the presence or absence of covert attention, respectively. These findings raise the possibility that, regardless of whether it is necessary, visual attention may play a determining role in visual appearance under certain conditions. However, because these previous studies involved physically defined stimuli, it remains unclear whether visual attention simply modulates pre-attentively computed structure as suggested by neurophysiological work (McMains & Kastner, 2011; Qiu et al., 2007), or whether structural computations depend on the state of attention. Rivalrous illusory figures are perfectly suited to address this issue: if attending to one illusory figure results in illusory contours that directly conflict with the form of another illusory figure, then structural computations must depend on attention.

To investigate the influence of voluntary attention on perceptual organisation, here we combined a novel illusory figure with an attentionally demanding task, exploiting human observers’ propensity to use illusory edges when making perceptual decisions (Gold, Murray, Bennett, & Sekuler, 2000). We developed a novel Kanizsa figure (**Fig. 1a**), in which “pacman” discs are arranged at the tips of an imaginary star. This figure includes multiple Gestalt cues that promote the segmentation and integration of various forms not defined by the physics of the stimulus (Harrison, Ayeni, & Bex, 2019). We predict that, because some of these cues suggest competing configurations, selective attention can bias which figure elements are assigned to figure and which to ground. Although such a hypothesis is relatively uncontroversial, the critical question is whether grouping via selective attention promotes illusory contour formation in direct conflict with competing implied form. For example, while the black inducers of **Figure 1a** form part of an implied star, in isolation the black inducers imply an illusory triangle that competes with both the star form as well as a second illusory triangle implied by the white inducers. The dependence of such perceptual organisation on voluntary attentional selection thus can reveal the extent of top-down processing on visual appearance. We therefore assessed whether the apparent organisation of the figure is determined by which inducers are attended.

**Figure 1.**
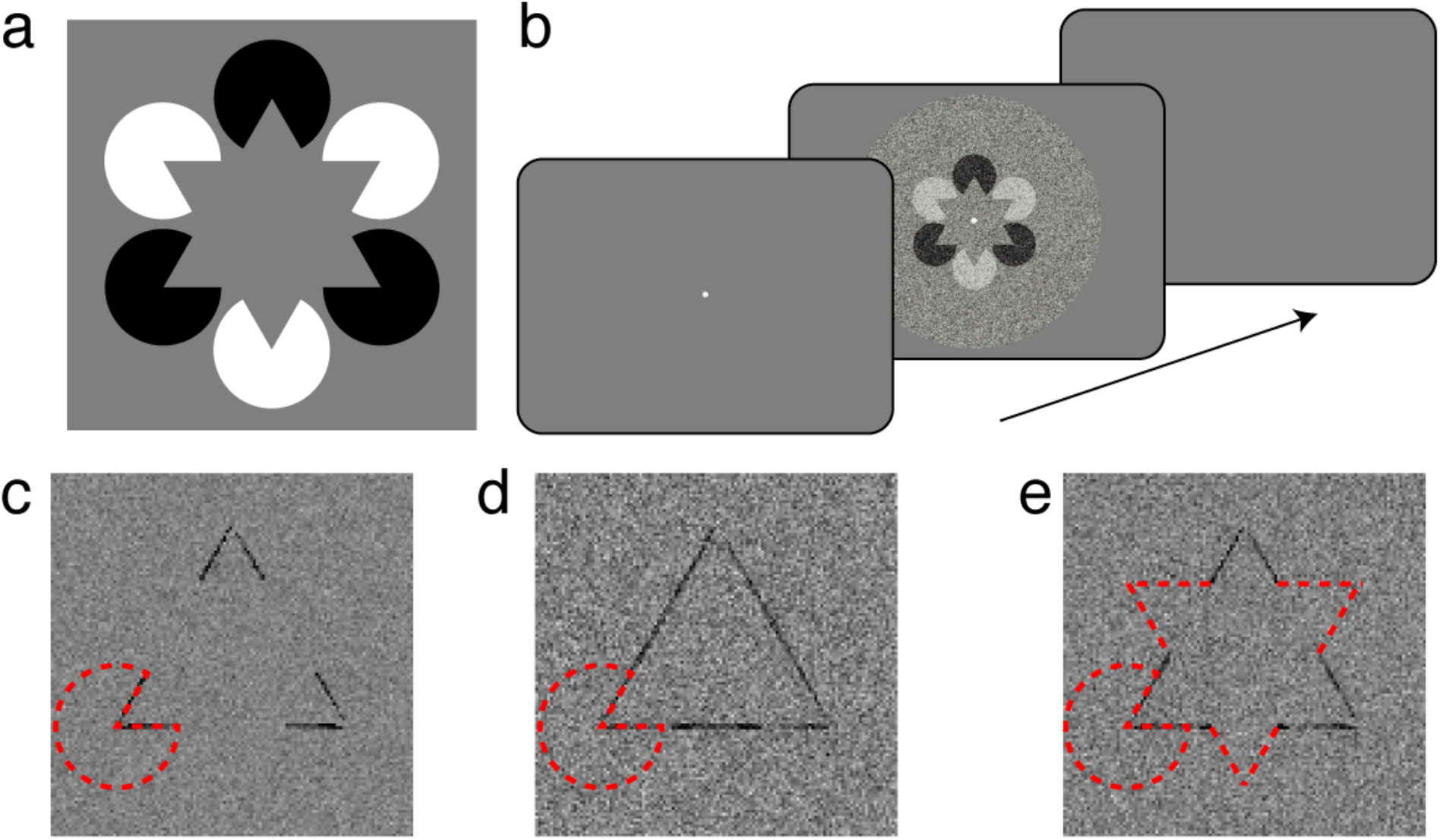
Novel illusory figure and design used to test the influence of attention on perceptual organisation. a) Our variant of the classic Kanizsa figure. “Pacman” inducers are arranged such that a star appears to occlude black and white discs. Whereas the ensemble of features may produce the appearance of a star, grouping features by polarity leads to competing illusory triangles. We test whether attending to one set of inducers (e.g. the white inducers) leads to interpolation of the illusory edge. b) Example trial sequence. After an observer fixates a spot, the illusory figure with overlaid Gaussian noise is displayed for 250ms. The observer’s task was to report whether the tips of the upright or inverted triangle were narrower or wider than an equilateral triangle. The target triangle was cued prior to, and held constant throughout, each testing block. The observer’s perceptual reports were then correlated with the noise on each trial to produce classification images. (c - e) Support vector machine (SVM) classifier images. We had a SVM classifier perform “narrow” vs “wide” triangle judgements after training it on three different protocols: (c) inducers, a (d) triangle, or a (e) star (see Methods). Dashed red lines show the location of a pacman for reference, and in (e) the tips of the star that do not influence classification.

## RESULTS

We used a response classification technique that allowed us to simultaneously assess where observers’ attention was allocated, and whether such attentional allocation resulted in visual interpolation of illusory edges. At the beginning of each block of testing, observers were cued to report the relative jaw size of the inducers forming an upright (or inverted) triangle, corresponding to the white (or black) elements in **Figure 1a**. By adding random visual noise to the target image on each trial (**Fig. 1b**), we could use reverse correlation to measure “classification images”. An observer’s classification image quantifies a correlation between each pixel in the image and the perceptual report revealing which spatial locations are used for perceptual decisions (Abbey, Eckstein, & Bochud, 1999; A. J. . J. Ahumada, Beard, & Ellis, 1997; A. J. Ahumada, 1996; A. Ahumada & Lovell, 1971; Beard & Ahumada, Jr., 1998; Gold et al., 2000).

We generated hypotheses regarding how observers’ voluntary attention may influence their perception of this figure. We used a support vector machine (SVM) classifier to judge small changes to a triangle image after training it on one of three different protocols (**Supp. Fig. 1**). First, we generated a prediction of the hypothesis that observers can attend to the correct inducers, but do not perceive illusory edges, by training a model to discriminate only the jaws of the inducers. This model is analogous to that of an ideal observer and reveals that only structure at the edges of the stimuli are used in generating a response (**Fig. 1c**). We next generated predictions of how illusory edges could be interpolated in this task. In one case, we assumed illusory contours would be formed between attended inducers. We thus trained the classifier to discriminate whether a triangle’s edges were bent outward or inward, and found a classification image approximating a triangle (**Fig. 1d**). In the other case, we assumed that, although selective attention may guide the correct perceptual decision, the illusory form of a star may be determined pre-attentively according to the physical structure of the entire stimulus. In this case, we trained the classifier to discriminate whether alternating tips of a star, i.e., the tips corresponding to a set of cued inducers, were relatively wide or narrow. The resulting classification image reveals edges that are interpolated beyond the inducers, but that they do not extend beyond the alternating star tips (**Fig. 1e**). These predictions not only provide qualitative comparisons for our empirical data, but they also allow us to formally test which training regime produces a classification image that most closely resembles human data.

To motivate observers to attend to only one possible configuration of the illusory figure, they were cued to report the relative jaw size (“narrow” or “wide”) of only a subset of pacmen positioned at the tips of an imaginary star (**Fig. 1a**). Each cued triangle was defined by three inducers, the jaw-sizes of which were varied from 60° (an implied equilateral triangle) according to an adaptive staircase (see Materials and Methods). In an early investigation into illusory contour perception, Ringach and Shapley (1996) found that observers’ perceptual thresholds were less than an angular degree with similarly sized stimuli. Specifically, observers were instructed to report only the jaw size of inducers forming an upward (or downward) triangle within a testing block. The non-cued inducer jaws varied independently of the cued inducers and thus added no information regarding the correct response. To derive the spatial structure used for perceptual decisions, we added Gaussian noise to each trial and classified each noise image according to the observers’ responses (**Fig. 1b**). To create the classification image for each observer, we summed all noise images for narrow reports and subtracted the sum of all noise images for wide reports (see Methods). We collapsed across inducer polarity by inverting the noise on trials in which the white inducers were cued, and across cue direction by flipping the noise on trials in which the downward facing illusory triangle was cued. The resulting images quantify the correlation between each stimulus pixel and the observer’s report. In order to analyse a single axis of emergent spatial structure, we first averaged each observer’s data with itself after rotating 120° and 240° such that correlations were averaged over the three sides of the triangle. Although this step involved bilinear interpolation of neighbouring pixels, no other averaging or smoothing was performed, and this averaging is therefore most likely to only reduce the strength of emergent illusory structure.

Classification images for three observers and their mean are shown in **Figure 2a** (see **Supp. Fig. 2a** for unrotated classification images). Images are normalised to the “attend upright black inducers” condition; black pixels indicate locations where dark and light noise was correlated with narrow and wide judgements, respectively, and white pixels indicate the opposite relationship. There are two obvious patterns that emerge. First, it is clear that observers based their reports on pixels within the jaws of the cued inducers, indicating that only some regions of the image – those aligned with the attended inducers – influenced perceptual decisions. Note the difference in the sign of the correlation between the edges and tips of the triangle – noise pixels in these regions have the opposite influence on narrow/wide decisions, which is likely due to an illusory widening of the jaw centre which is not registered by the SVM (cf. **Fig. 1d**). Second, the edges clearly extend beyond the red inducer outline shown in the mean image, revealing observers’ reports were influenced by illusory contours. However, it is also apparent that the spatial structure is non-uniform, with weaker correlations in the centre of the illusory edges than in the corners of the inducers. We therefore quantitatively test the extent of illusory contour formation below.

**Figure 2.**
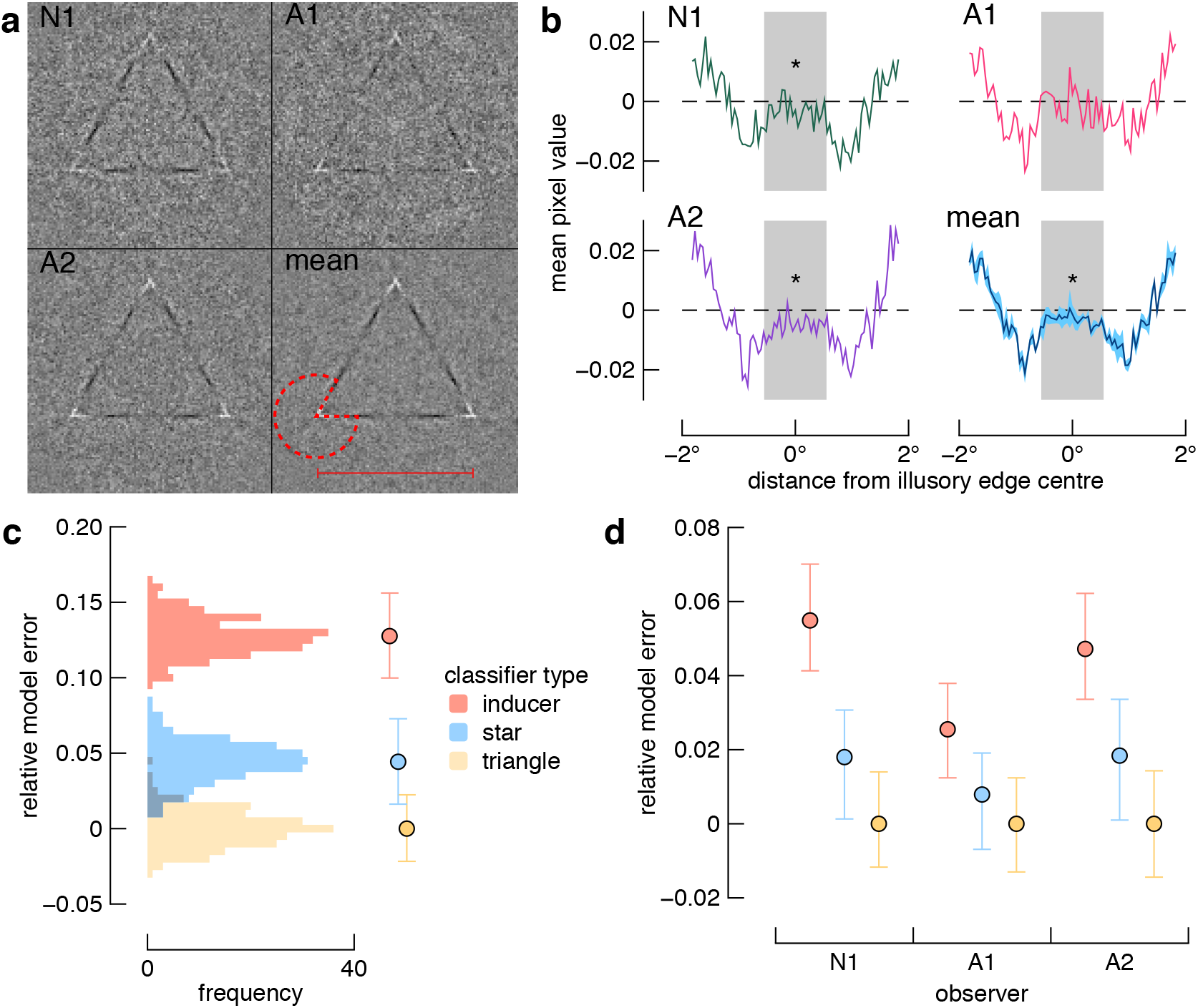
Classification image results. a) The individual and average classification images, normalized to the “attend upright black inducers” condition. Black pixels indicate locations where dark and light noise was correlated with narrow and wide judgements, respectively, and white pixels indicate the opposite relationship, after 9984 trials per participant. Data have not been smoothed, but were first averaged across triangle edges and cropped to be 122×122 pixels. In the mean image, a pacman outline is shown for reference, and a red line indicates the spatial range of the implied triangle edge (from which data in (b) are shown). b) Pixel values along the illusory edge. The grey shaded region corresponds to a conservative estimate of the extent of a gap in the edge that would appear if observers necessarily saw a star shape (e.g. Fig. 1e). The blue shaded region in the mean plot shows ±one standard error; asterisks indicate differences from zero (BF_10_ > 10 and p < 0.05; see text). N1 is the naïve participant; A1 and A2 are authors. c) Comparison of SVM models for averaged data. Distributions show comparisons of the mean classification image to the output of each SVM prediction, repeated 200 times. Data points and error bars represent the mean and 95% confidence intervals, respectively, for each SVM training regime. Model error has been normalised relative to the model with the least error, which is the model in which the SVM is trained to perceive a triangle within the attended inducers. d) Comparison of SVM models for each observer. Colours are as per panel (c).

To test whether the illusory edge interpolation extended into the region of the implied competing figure, we performed two analyses. First, we used Bayesian and Students’ one-sampled t-tests to assess the pixel values along the edge of the triangle implied by the attended inducers (see red line in **Fig. 2a**). We selected only pixels that fell within the bounds of the competing implied triangle (see Methods and grey shaded regions of **Fig. 2b**), and found that these 18 pixels were below zero for the naïve participant (mean and sem: −3 ± .9 × 10^−3^, BF_10_=18.365, t(17)=3.585, p=0.002, d = 0.845), observer A2 (mean and sem: −5 ± .7 × 10^−3^, BF_10_=8,141.356, t(17)=6.944, p<0.001, d = 1.637), and the group (mean and sem: − 3 ± .4 × 10^−3^, BF_10_=16,580, t(17) = 7.38, p<0.001, d = 1.738), but not for A1 (mean and sem: −1 ± 1 × 10^−3^, BF_10_=0.431, t(17)=1.15, p=0.266, d = 0.204). The lack of a difference in observer A1 may be due to a difference in task related strategy and or increased lapse rate.

We next quantified the spatial structure content of the classification image by testing which prediction generated by the SVM was most similar to the human data (see **Fig. 1c-e**). For each model, we generated 200 predictions, each with a unique distribution of noise, and computed the sum of squared errors between predictions and the mean classification image produced by the human observers (see Materials and Methods). The resulting distributions of error, normalised to the best model, are shown in **Figure 2c**, and reveal that the model in which we trained the classifier to perceive a complete triangle is the best fit to the data (z-test comparing the mean error for the triangle SVM versus the distribution of error for the star or inducer SVM: p’s < 0.0001). The pattern of results was the same for all observers (**Fig. 2d**): for the inducer, star, and triangle templates, respectively, the mean standardised model errors (±one standard deviation) were N1: 0.055 (0.007), 0.018 (0.008), 0 (0.007); A1: 0.026 (0.006), 0.008 (0.006), 0 (0.007); A2: 0.047 (0.008), 0.018 (0.008), 0 (0.007). Z-tests comparing the mean error for the triangle SVM versus the distribution of error for the star or inducer SVM were all significant (all p’s < 0.0001). Taken together, these analyses reveal illusory contour formation between attended visual elements, and this interpolation occurred despite the contour conflicting with equally plausible implied spatial structure.

We next tested the spatial specificity of illusory contour formation. In the preceding analyses presented in Figure 2b, we selectively tested only a single row of pixels aligned with the mouths of the inducers. For the two participants who showed a clear effect, we next tested how spatially specific visual interpolation was by repeating the same analysis but for the row of pixels above and below the triangle boundary implied by the geometry of the attended inducers. Quite surprisingly, we found good evidence that there was an absence of illusory contour formation for the pixels below the implied triangle boundary (N1: BF_01_ = 3.19; A2: BF_01_ = 3.31), and equivocal evidence for the pixels above the implied triangle boundary (N1: BF_10_ = 1.05; A2: BF_01_ = 1.83). We therefore found evidence that only a single row of pixels extending between the inducer edges contributed to observers’ perceptual decisions. These results thus reveal that the strength of illusory contours was highly precisely aligned to the geometry of the triangle implied by the attended inducers. Consistent with this observation, psychophysical thresholds for identifying the relative inducer jaw size were reliably highly precise across testing sessions (see **Supp. Fig. 2b**). Across sessions, the mean thresholds (±one standard error) for observer N1, A1, and A2, were 0.86° ± .03°, 0.84° ± .02°, and 0.66° ± .03°.

Our data further address the extent to which the non-cued figural elements may have influenced perceptual judgements. In our experiment, the non-cued inducer jaw size was independent of the cued inducer jaw size, and was thus uninformative of the correct report. Indeed, we found no evidence in the classification image that observers’ perceptual decisions were guided by these task-irrelevant cues. We modelled the possibility that these non-cued elements were nonetheless grouped to form a star. In such a case where a star was perceived, the task could still be performed accurately were observers to base their reports on only the edges shared by the star and the triangle implied by the cued inducers. As expected, the SVM prediction of pre-attentive figure-ground segmentation shows gaps in the sides of the classification image triangle (**Fig 1e**). Note that this model is equivalent to observers having perceived a whole star, but with a later stage attentional signal focussed on only some regions of the pre-computed figure. Because we designed our illusory figure to be geometrically invertible, the extent of the illusory star form is pronounced if we sum the model’s classification image with a flipped version of itself (**Fig. 3a**). In **Figure 3b**, we show the result of performing this step with the observers’ average classification image. Very similar patterns of results were found for all individual images (**Supp. Fig. 3**).

**Figure 3.**
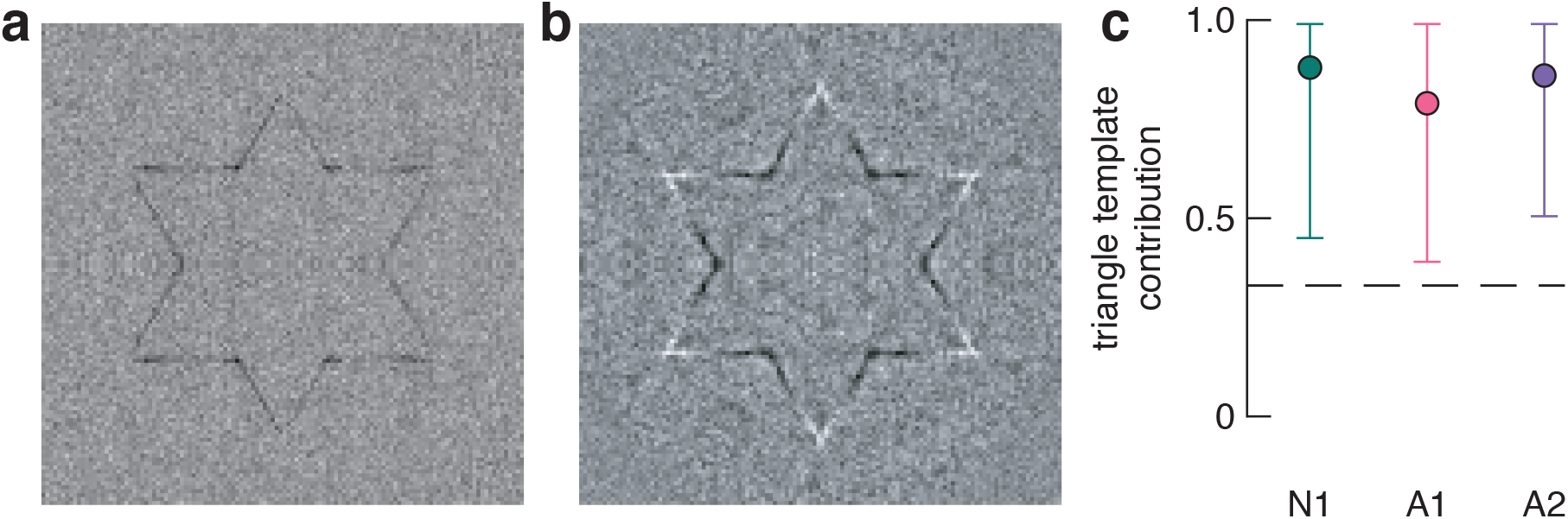
Pre-attentive grouping. a) Geometric form prediction of unattended grouping. The classification image derived from our SVM was summed with a flipped version of itself. Note that the inner corners of the star are well aligned due to the design of our original Kanizsa figure. b) Geometric form in observers’ data. The mean classification image was summed with a flipped version of itself, and reveals the strength of illusory edges are well aligned to the implied star. c) Results of mixture modelling used to explore the correspondence between fluctuations of illusory edge strength and implied figure geometry. The best fitting model for each observer was one in which attention determined perceptual outcome in 84% of trials. The dashed line indicates the proportion expected from a purely stochastic process. Error bars show 95% confidence intervals.

Although flipping the classification image and producing a star-like figure may be somewhat trivial, more important is how the edges of the star are formed. We wish to test whether the extent of the illusory lines matches what would be expected were observers to have relied on a pre-attentively computed star form, rather than two superimposed triangles determined by attentional selection. The critical aspect of the star-like figure shown in Figure 3b is therefore whether or not the lines that form the star have terminators at the point where they intersect (ie. the inner corners of the implied star). This is the case in Figure 3a because we are using the star model to generate the classification image. For observers’ data in Figure 3b, on first glance the same appears to be the case: the emergent features appear to stop precisely at the point of intersection, suggesting observers perceived a star but based their reports on only some parts of this figure. These qualitative results, however, are in contrast to the SVM analysis, presented above (Fig. 2), in which we found that correlated noise in observers’ classification images is best explained by observers having attended to the cued triangle. Thus, a remaining critical question, which we address below, is whether we can quantify the proportion of trials in which observers relied on different forms implied by the inducers.

In contrast to our initial quantitative analyses, the results of which suggest visual attention determines which of two illusory triangles were perceived on a trial (**Fig. 2**), qualitative inspection of the star-like form shown in Figure 3b suggests observers may have based their reports on only some parts of a pre-attentively computed star. However, there are at least three possible explanations for the near-perfect alignment of changes in illusory edge strength with the implied star figure (**Fig. 3b**). First, as discussed, a similar classification image would have been obtained had observers perceived a star on every trial, a possibility which we discounted by our quantitative analysis of the illusory edge, described above (Fig. 2). Second, this qualitative result could be generated if trial-by-trial perceptual organisation was stochastic, such that observers perceived each possible configuration approximately equally often across trials. Under this hypothesis, the resulting illusory contours shown in **Figure 2a** are incidental rather than being determined by observers’ attentional goals. The third possible explanation is that observers’ voluntary allocation of attention determined the outcome on most, but not all, trials. To distinguish between the two latter possibilities, we used mixture modelling to quantify the proportion of trials in which observers’ percept depended on attentional instructions (see Materials and Methods). A purely stochastic process would be implied were the proportion of trials accounted for by the triangle template no different from 0.33 (i.e. the apparent top-most surface was equally often a star, the cued triangle, or the non-cued triangle, see **Fig. 1c-e**). However, in the best fitting model, the attention-contingent triangle template contributed to 84% of trials on average, which is much greater than expected by a stochastic process (**Fig. 3c**). At the individual level, the triangle contributions (and 95% confidence intervals) for N1, A1, and A2 were 88% (45% - 100%), 79% (39% - 99.5%), and 86% (50.5% - 100%), respectively. This mixture modelling is thus consistent with observers’ attentional goals determining illusory contour interpolation on the vast majority of trials.

## DISCUSSION

We used classification images to address whether voluntary attention determines a scene’s apparent visual structure. Using a psychophysical response classification paradigm we tested which of three competing model predictions best describes the influence of attention on illusory contour formation. Our results clearly show that voluntary attention can guide the fundamental processes involved in perceptual organisation of illusory structure.

Unlike previous studies that show visual attention modulates the appearance of physically defined surfaces (e.g., attending to different surfaces of the Necker cube (Necker, 1832)), our study shows a rich interaction between attention and endogenously generated percepts. Classification images reveal the spatial location of noise elements that influence observers’ responses, whereas interpreting subjective phenomenology is more difficult. However, our stimulus design ensured that the classification images reveal information about the perceived depth order of image elements. The presence of lines in the classification image that extend between the inducers is clear evidence that at least two of three observers based their judgements on the perception of a figure whose edges occlude the competing (non-cued) shape information. Given that the illusory edges of the triangle implied by the attended inducers directly conflict with the regions of the competing implied figures (i.e., the star and inverted triangle), our finding that illusory edges were interpolated between attended inducers reveals that attention can determine depth order, even when figures and ground are illusory. Spatial structure is thus computed by neural operations that are at least partially contingent on the voluntary state of the observer. The precision of illusory contours was nonetheless tightly aligned to the geometry of luminance defined structure, indicating these inferential processes are also highly contingent on scene or task context. Indeed, observers’ psychophysical thresholds for the inducer task reveal a correspondence between their precise objective psychophysical performance and subjective classification image.

We found clear inter-participant differences in the classification images. First, we found a clear effect of edge completion in our initial analysis of edge completion in only two out of three observers (Fig. 2). Such a difference across participants is not exceptional: Gold and Shubel (2006) also found classification image evidence of illusory edges in two out of three participants. Nonetheless, although the effect did not reach significance for one observer in our data, the same general direction of results was found in both the classification image analysis (Fig. 2b) and the same results were found in the individual SVM model comparisons (Fig. 2d). A degree of homogeneity of our results across participants is also reflected by the fact that the group-average effect was significant. Importantly, the critical effect of a fully interpolated illusory edge was found in the naïve observer’s data, and, across participants, we found relatively strong effect sizes of d = 0.845 (N1), d = 1.637 (A2), and d = 0.204 (A1) despite not being significant for A1. The second inter-participant differences we found were in the raw classification images that reveal varying degrees of completeness (Supp. Fig. 2). For the two observers where the effect was significant, at least two edges of the triangle are clearly visible, and for the remaining observer one edge is clearly visible. We can think of at least three possible explanations for these individual differences (similar between-observer differences were reported by Gold et al, 2000, and Gold & Shubel, 2006). First, observers may have interpolated the edge of a single or pair of unconnected lines between the cued inducers. Second, observers perceived a triangle, but only used part of this triangle to perform the task. Third, there are individual biases in attentional allocation that differentially influenced interpolation of the different edges. Given the strength of the Kanizsa illusion, i.e., the perception of a triangle, we think that the latter two explanations are more likely, however we cannot definitively show this with the current data. The conclusion that attention influences illusory contour formation is equally valid under either of these explanations.

We were able to quantify the influence of non-cued stimuli on perception by measuring a classification image across the entire stimulus. We found that changes in the strength of illusory contour formation between attended inducers were aligned with form implied by the non-cued inducers. Our mixture modelling suggests that the non-cued stimuli influenced performance on approximately 16% of trials. Such a contribution of task-irrelevant features on perceptual decisions could be attributed to lapses in attentional allocation, or variability in the feed-forward processing of the incoming signal. Measuring perceived form in the absence of visual attention is notoriously difficult (Wagemans et al., 2012), which is perhaps one reason why many studies of figure-ground organisation rely on single-unit recordings. Whereas neurophysiological recordings have revealed the brain regions involved in perceptual organisation, they have left open the question of perceptual phenomena. Our data show that the influence of attention on perception is constrained by task-irrelevant information, providing yet further evidence that visual experience is the combination of both bottom-up and top-down processes. This conclusion sheds light on previous work in which competing colour adaptation after-effects are biased according to alternating illusory contours at a similar location (van Lier, Vergeer, & Anstis, 2009). In these demonstrations, the onset of inducer elements likely attracts an observer’s attention, resulting in perceptual completion processes specific to only the implied shape of attended elements. Surface filling-in would then follow the contours of the implied form (Poort et al., 2012). Indeed, other recent research from our lab reveals similar interactions may occur between attention and surface filling-in (Harrison et al., 2019).

The influence of attention on figure-ground segmentation may be explained by feedback signals from the lateral occipital complex (Murray et al., 2002; Stanley & Rubin, 2003) that could act as early as V1 (Wannig et al., 2011), but also may involve modulating responses of border-ownership cells in V2 (Qiu et al., 2007). Border-ownership cells indicate which side of a border is an object versus ground. Previous work showing the activity of border-ownership cells is modulated by visual attention (Qiu et al., 2007) has been limited to luminance-defined borders. Our finding that information inferred by the visual system is influenced by voluntary attention suggests that attentional modulation of border-ownership may similarly apply to illusory contours (R von der Heydt et al., 1984). Early psychophysical work suggested that illusory contours are perceived in the absence of attention (Davis & Driver, 1994; Mattingley, Davis, & Driver, 1997), but did not address the question of whether illusory contours can be formed *because* of voluntary attention, which we have shown here. Our findings are also distinct from other recent work that found attention can influence the appearance of existing surfaces (Tse, 2005). In our study, visual attention had a causal role in forming the structure from which perceptual decisions were made. We anticipate that our simple stimulus and task design may prove to be a useful neurophysiological assay to test further the neural substrates governing the interaction between voluntary attention and perceptual organisation.

## MATERIALS AND METHODS

### Observers

Three healthy subjects, one naïve (N1) and two authors (A1 & A2 corresponding to authors RR and WH, respectively), gave their informed written consent to participate in the project, which was approved by the University of Cambridge Psychology Research Ethics Committee. All procedures were in accordance with approved guidelines. Simulations were run to determine an appropriate number of trials per participant to ensure sufficient statistical power, and our total sample is similar to those generally employed for classification images. All participants had normal vision.

### Apparatus

Stimuli were generated in MATLAB (The MathWorks, Inc., Matick, MA) using Psychophysics Toolbox extensions (Brainard, 1997; Cornelissen, Peters, & Palmer, 2002; Pelli, 1997). Stimuli were presented on a calibrated ASUS LCD monitor (120Hz, 1920×1200). The viewing distance was 57 cm and participants’ head position was stabilized using a head and chin rest (43 pixels per degree of visual angle). Eye movement was recorded at 500Hz using an EyeLink 1000 (SR Research Ltd., Ontario, Canada).

### Stimuli and task

The stimulus was a modified version of the classic Kanisza triangle. Six pacman discs (radius = 1°) were arranged at the tips of an imaginary star centred on a fixation spot. The six tips of the star were equally spaced, and the distance from the centre of the star to the centre of each pacman was 2.1°. The fixation spot was a white circle (0.1° diameter) and a black cross hair (stroke width = 1 pixel). The stimulus was presented on a grey background (77.5 cd/m^2^). The polarity of the inducers with respect to the background alternated across star tips. For half the trials, the three inducers forming an upright triangle were white, while the others were black, and for half the trials this was reversed. Inducers had a Weber contrast of .75.

We added Gaussian noise to the stimulus on each trial to measure classification images. Noise was 250 × 250 independently drawn luminance values with a mean of 0 and standard deviation of 1. Each noise image was scaled without interpolation to occupy 500 × 500 pixels, such that each randomly drawn luminance value occupied 2 × 2 pixels (.05° × .05°). The amplitude of these luminance values was then scaled to have an effective contrast of 0.125 on the display background, and were then added to the Kanizsa figure. Finally, a circular aperture was applied to the noise to ensure the edges of the inducers were equally spaced from the noise edge (**Fig. 1b**).

The jaw size of inducers was manipulated such that they were wider or narrower than an equilateral triangle, which would have exactly 60° of jaw angle for all inducers. The observer’s task was to indicate whether the jaws of the attended inducers was consistent with a triangle that was narrower or wider than an equilateral triangle. Prior to the first trial of a block, a message on the screen indicated which set of inducers framed the “target” triangle, and this was held constant within a block but alternated across blocks. The polarity of the target inducers and whether the triangles were narrow or wide was pseudorandomly assigned across trials such that an equal number of all trial types were included in each block. The relative jaw size of attended inducers was independent of the unattended inducers; thus, the identity of the non-target triangle was uncorrelated with the correct response.

Each trial began with the onset of the fixation spot and a check of fixation compliance for 250 ms. Following an additional random interval (0-500 ms uniformly distributed), the stimulus was presented for 250 ms, after which only the background was presented while observers were given unlimited duration to report the jaw size using a button press. The next trial would immediately follow a response. Throughout the experiment, eye tracking was used to ensure observers did not break fixation during stimulus presentation. If gaze position strayed from fixation by more than 2° the trial was aborted and a message was presented instructing them to maintain fixation during stimulus presentation, and then the trial was repeated. Such breaks in fixation were extremely rare for all participants.

A three-down one-up staircase procedure was used to progress the difficulty of the task by varying the difference of the jaw size from 60° (i.e., from what would form an equilateral triangle). On each trial an additional angle was randomly added or subtracted to the standard 60° inducers. The initial difference was 2°. Following three correct responses, this difference would decrease by a step size of 0.5°, or would increase by the same amount following a single error. When an incorrect response was followed by three correct responses (i.e., a reversal), the step size halved. If two incorrect responses were made in a row, the step size would double. If the step size fell below 0.05°, it would be reset to 0.2°. Blocks consisted of 624 trials which took approximately 20 minutes including a forced break. Each observer completed 16 blocks for a total of 9984 trials, which took a total of approximately five hours duration spread over multiple days and testing sessions. To familiarize observers with the task, they underwent two training blocks of 624 trials each with no noise. They then were shown the stimulus with noise, and completed as many trials as they felt was required before starting the experimental blocks.

### Support vector machine models

Support vector machine (SVM) classifiers were trained and tested in MATLAB. We generated (3) hypotheses by training SVM classifiers on images of the *i*) inducers, *ii*) a triangle, or *iii*) a star. We trained the classifiers using a quadratic kernel function and a least squares method of hyperplane separation. The training images consisted of two exemplars (“narrow” and “wide”) with no noise (**Supp. Fig. 1**). To generate hypotheses in the form of classification images, we used each of the classifiers to perform narrow/wide triangle judgements (trials = 9984), with an equilateral triangle; thus, classification was exclusively influenced by the noise in the image.

### Data and statistical analysis

The 9984 noise images for a participant were separated according to perceptual report (“narrow” or “wide”). To collapse across inducer polarity, we inverted the sign of noise on trials in which the cued inducers were white. We also collapsed across upright and inverted cue conditions by spatially flipping the noise on inverted trials. To calculate which spatial locations influenced perceptual reports, we used a standard classification analysis in which each trial is classified according to the observer’s response with respect to the stimulus shown on that trial (Gold et al., 2000; Gold & Shubel, 2006; Mareschal, Dakin, & Bex, 2006; Neri & Heeger, 2002). Each stimulus was either narrow or wide (*S_narrow_* or *S_wide_*), and each response was either narrow or wide (*R_narrow_* or *R_wide_*), giving four trial types: 1) *S_narrow_ R_narrow_*, 2) *S_wide_ R_narrow_*, 3) *S_wide_ R_wide_*, 4) *S_narrow_ R_wide_*. The classification images were generated by averaging and combining these response types according to the equation:

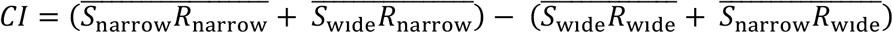

The resulting classification image shows the strength of correlation between each pixel’s location with the perceptual report made by the observer. Images are normalised to the “attend upright black inducers” condition, such that black pixels indicate locations where dark luminance noise was correlated with a narrow response, and light luminance noise was correlated with a wide response. Conversely, white pixels indicate locations at which light luminance noise was correlated with a narrow response, and dark luminance noise was correlated with a wide response. The polarity and intensity of a given location thus provides information regarding that location’s contribution to perceptual decision making. To average across emergent triangle edges, we further summed the image with itself two times after rotating 120° and 240° using Matlab’s “imrotate” function using bilinear interpolation. This procedure results in a classification image that is invariant across edges such that analysis of one edge summarises all three edges. Note that this is a conservative estimate of the classification image and any spurious structure will only be diminished. To test for correlated pixels along the illusory edge of the classification image, we extracted 18 pixels along the bottom edge of the implied triangle, but within the bounds of the implied star tip (see bottom right panel of **Fig. 2a**). To ensure that these pixels were not contaminated by averaging of nearest-neighbour pixels during rotation, described above, we excluded the three pixels closest to the inner corners of the star. We conducted a one-sample, two-tailed Bayesian and Students’ T-Test on these pixel values using JASP software (JASP Team, 2017). Reported effect sizes are Cohen’s d.

We performed the model comparisons in **Figure 2c** by first normalising the noise of the mean classification image and each SVM prediction such that the sum of squared error of each image equalled 1. We then subtracted the mean classification image from each prediction, and found the sum of squared error of the resulting difference. Finally, we normalised the difference scores to the model with the least error by subtracting from each distribution the mean of the distribution with the lowest error. This process was repeated for 200 repetitions of each SVM prediction. The mixture modelling (**Fig. 3c**) was performed similarly, but we further used Monte Carlo simulations to estimate the proportion of trials in which a triangle was perceived. In this case, each set of 200 simulated experiments included a proportion of triangle template trials, ranging from 0.33 (chance) to 1. We validated this model fitting procedure by generating a simulated classification image with a known generative template, or with proportional mixtures of templates, and then verified the model fitting returned results that approximated the ground truth. The Monte Carlo simulations were highly accurate for a range of simulated proportions, but slightly overestimated the contribution of the triangle template when the ground truth contribution was close to 0.33, and, conversely, slightly underestimated its contribution when the triangle was the only contributor.

## Data availability

The data that support the findings of this study are available from the corresponding author upon request.

## ACKNOWLEDGEMENTS

We are indebted to Peter Bex who developed the novel Kanizsa figure with us and provided helpful feedback on our study design and results. We also thank Tom Wallis for feedback on an earlier draft which led to the mixture modelling and overall improvements in the manuscript. This research was supported by funding to W.J.H. from King’s College Cambridge and the National Health and Medical Research Council of Australia (APP1091257).

## AUTHOR CONTRIBUTIONS

Both authors designed the experiment and collected the data. WJH analysed the experimental data, RR performed the SVM analyses, and both authors performed the model comparisons. Both authors contributed equally to the writing of the manuscript.

We declare we have no competing interests.

---

**Supplementary Figure 1.**
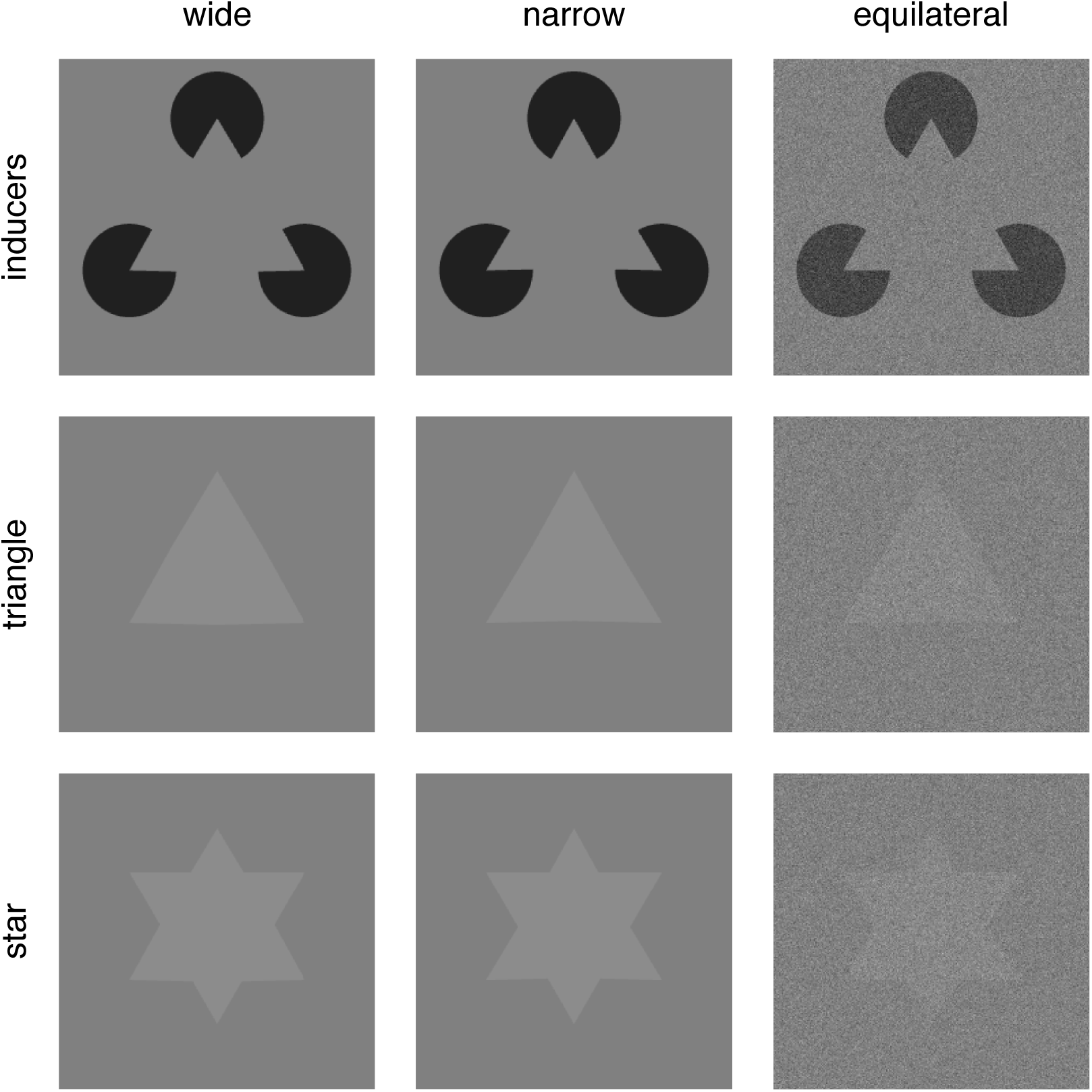
Support vector machine (SVM) images. From left to right, the first two columns show examples of wide and narrow exemplar images used to train the SVM in the inducers, triangle, and star protocols. The column on the right shows examples of the test image for each protocol.

**Supplementary Figure 2.**
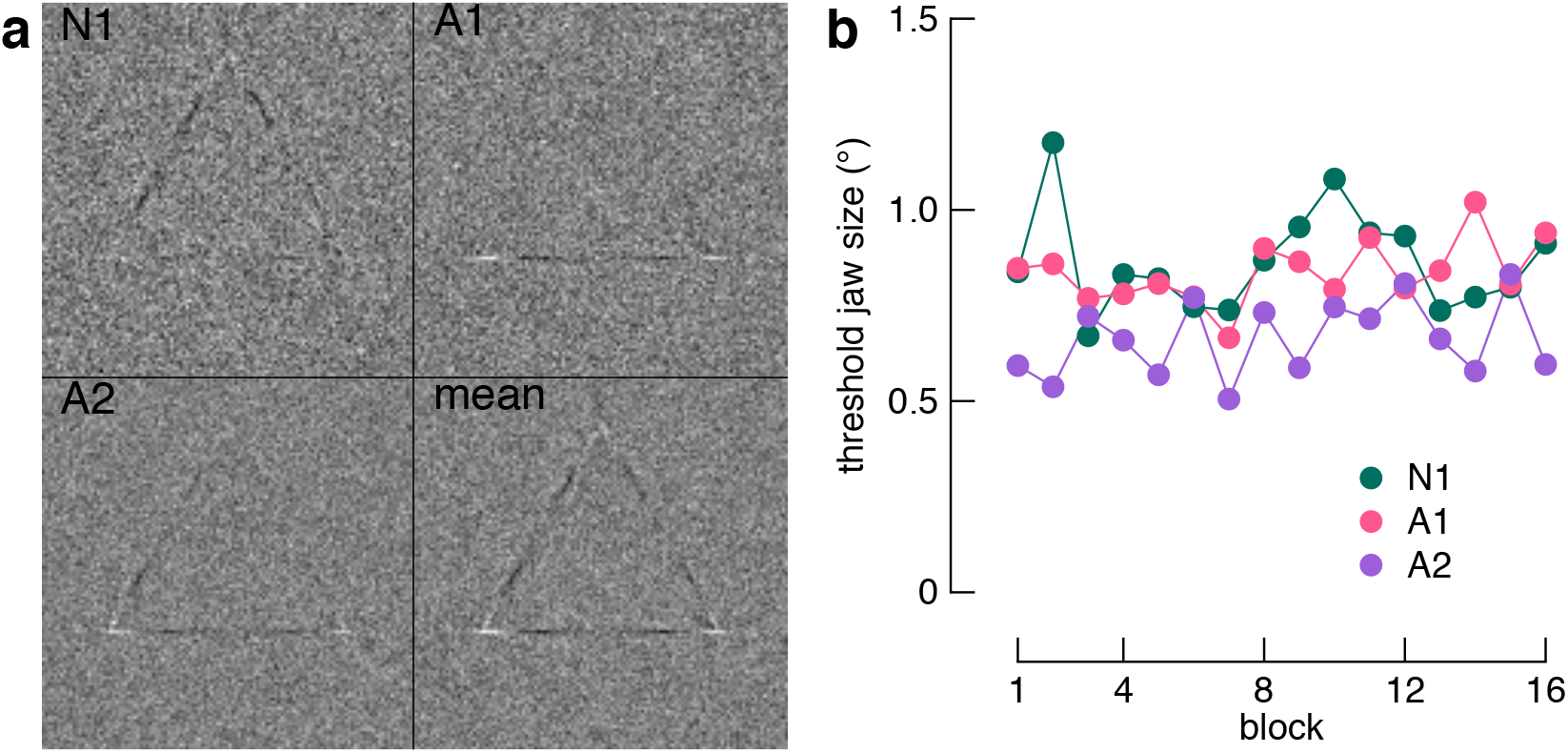
Raw classification images and psychophysical performance. a) Classification images without averaging of edges via rotation. Note that the individual images show varying degrees of a complete triangle. One explanation for this is that observers perceived a partial shape, e.g., a single or pair of unconnected lines between the cued inducers. Given the strength of the Kanizsa illusion in producing the percept of a triangle, rather than a partial shape, a more likely explanation is that observers perceived a triangle, but only used part of this triangle to perform the task. b) Threshold performance across blocks shown separately for each observer. Thresholds were the midpoint of a cumulative Gaussian fit to accuracy data for each session.

**Supplementary Figure 3.**
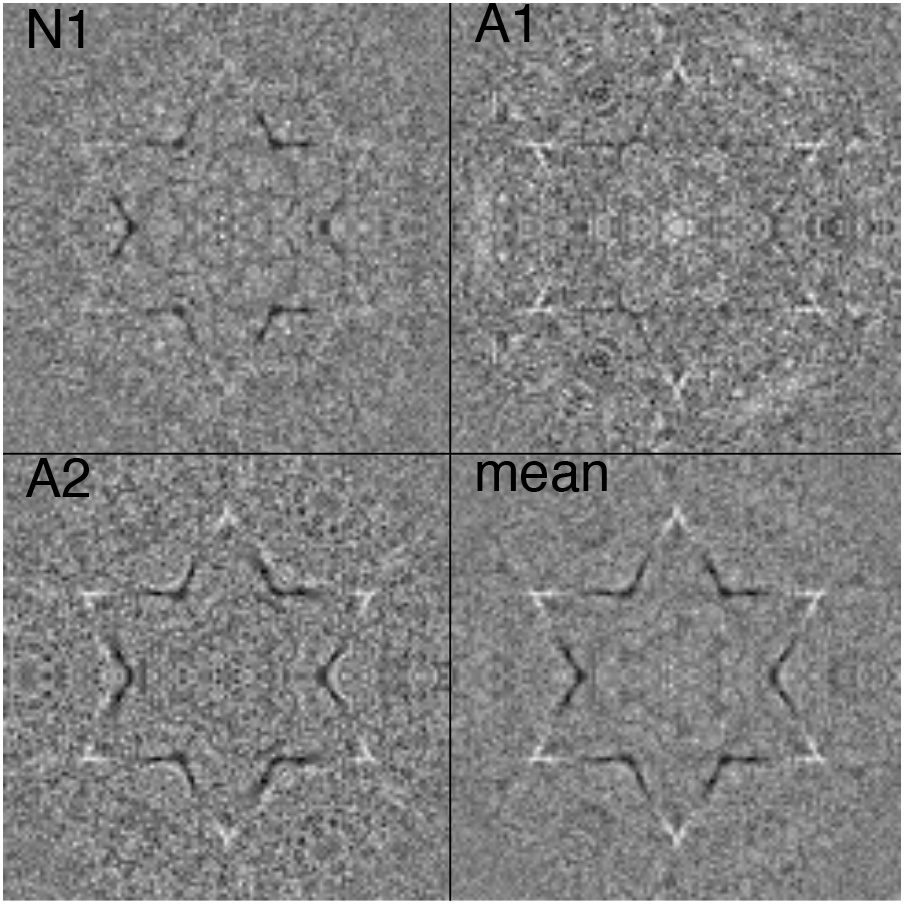
Individual classification images revealing a potential influence of non-cued structure. These images were created by summing each classification image with a flipped version of itself. Note that the emergent structure aligns to the geometry of the star implied by our Kanizsa figure (**Fig. 1a**).

